# Protein Function Prediction with Pretrained ProtT5 Embeddings and Gradient Boosting

**DOI:** 10.64898/2026.04.27.721184

**Authors:** Jett Appel, Nathan Butcher

## Abstract

Protein function prediction remains a central challenge in computational biology due to the extreme sparsity and long-tail distribution of Gene Ontology (GO) [1] annotations. Advances in protein language models enable the extraction of dense, fixed-length representations from amino acid sequences, offering a scalable alternative to hand-picked features such as physicochemical properties. In this work, we evaluate a transformer-based embedding approach using ProtT5-XL combined with classical and modern multi-label classifiers for Gene Ontology prediction in the CAFA-6 setting. Fixed-length embeddings were generated via mean pooling of transformer hidden states and used as input to one-vs-rest logistic regression, gradient-boosted decision trees, and a neural network. Models were evaluated on held-out validation data with a focus on threshold selection, prediction sparsity, and behavior across frequent and rare GO terms. Gradient boosting consistently provided the best balance between predictive performance and stable prediction behavior, motivating its use for ontology-specific predictors across molecular function, biological process, and cellular component annotations. This study highlights practical modeling choices for large-scale protein function prediction using pretrained sequence embeddings and provides an interpretable baseline for future CAFA evaluations.

## 1 Introduction

Accurately annotating protein function is essential for understanding biological systems, yet experimental characterization remains costly and incomplete. As a result, large public databases such as the Gene Ontology (GO) contain annotations that are both incomplete and highly imbalanced, with a small number of common terms and a long tail of rare functions supported by only a handful of proteins. This extreme sparsity poses a major challenge for supervised learning approaches to protein function prediction.

Recent progress in protein language models [2][3] has transformed sequence-based modeling by enabling the extraction of contextual representations directly from amino acid sequences. Models such as ProtT5 [3], trained on tens of millions of unlabeled protein sequences, capture structural and functional information without requiring multiple sequence alignments or handcrafted features. These embeddings provide a natural foundation for scalable downstream prediction tasks, including multi-label GO annotation.

Despite their promise, practical questions remain regarding how best to combine pretrained embeddings with downstream classifiers in highly sparse, multi-label settings. In particular, the choice of classifier, probability thresholding strategy, and handling of rare GO terms can substantially affect prediction behavior and evaluation metrics. Additionally, different GO ontologies, including molecular function (MF), biological process (BP), and cellular component (CC), exhibit distinct label distributions and semantics, suggesting that a single unified model may be suboptimal.

In this work, we investigate a straightforward and interpretable pipeline for protein function prediction using ProtT5-derived embeddings and ontology-specific classifiers within the CAFA-6 framework [4]. We compare linear, tree-based, and neural network models on held-out validation data, analyze prediction sparsity and threshold sensitivity, and motivate the use of gradient-boosted decision trees for final ontology-specific predictors. Rather than optimizing solely for leaderboard performance, our goal is to characterize modeling tradeoffs and provide a reproducible baseline for large-scale protein function annotation using pretrained sequence representations.

## 2 Methods

### 2.1 Protein embeddings using a pretrained transformer model

Protein amino acid sequences were obtained from the CAFA-6 training and test FASTA files. Each sequence was parsed and associated with a unique identifier (EntryID). For compatibility, whitespace was inserted between each amino acid character prior to tokenization.

ProtT5-XL-Uniref50, a transformer protein language model, was used to obtain fixed-length numerical representations to allow for downstream multi-label classification. The model was trained on the UniRef50 dataset consisting of approximately 45 million protein sequences and contains approximately 3 billion parameters.

The embeddings were generated using the Hugging Face implementation [5]. The model was loaded in evaluation mode, and all parameters were frozen during inference. For each protein sequence, token embeddings were computed using a forward pass through the transformer encoder. Embeddings were generated using mean pooling across the sequence length dimension of the final hidden layer, resulting in a 1024-dimensional embedding vector for each protein.

For a given protein sequence with token-level hidden states, the protein embedding was computed as:

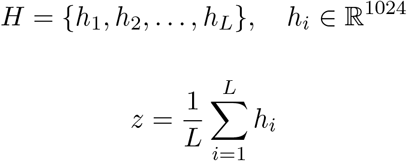

The embeddings were generated in batches using two NVIDIA Tesla T4 GPUs [7] due to the computational and memory demands of the ProtT5-XL model. A batch size of 4 was used, and embeddings were computed sequentially across multiple sessions for both the training and testing data. The final output consisted of one fixed-length 1024-dimensional embedding vector per protein sequence. These vectors were merged with protein identifiers and used as input features for multi-label classification modeling across molecular function, biological process, and cellular component ontologies.

### 2.2 Dataset preprocessing and model preliminary analysis

The CAFA-6 training set was used, which consisted of protein sequences annotated with GO terms. These terms were grouped per protein and split by ontology aspect. For this preliminary analysis, the initial focus was on the MF ontology.

Protein embeddings were obtained from the pretrained ProtT5 transformer model. These embeddings were merged with MF GO annotations using protein accession identifiers.

The predictions were restricted to the top 500 most frequent MF GO terms in the training data since the MF annotations are highly sparse and right-skewed. This strategy balances biological coverage and computational efficiency while avoiding labels with extreme sparsity that can cause model instability.

The machine learning (ML) workflow is detailed in Figure 1, where out of 58,001 proteins, 80% was split into training and 20% into validation using a fixed random seed to ensure reproducibility. To enable rapid preliminary model comparison, 8,000 proteins were randomly subsampled from the training split using a fixed seed.

**Figure 1:**
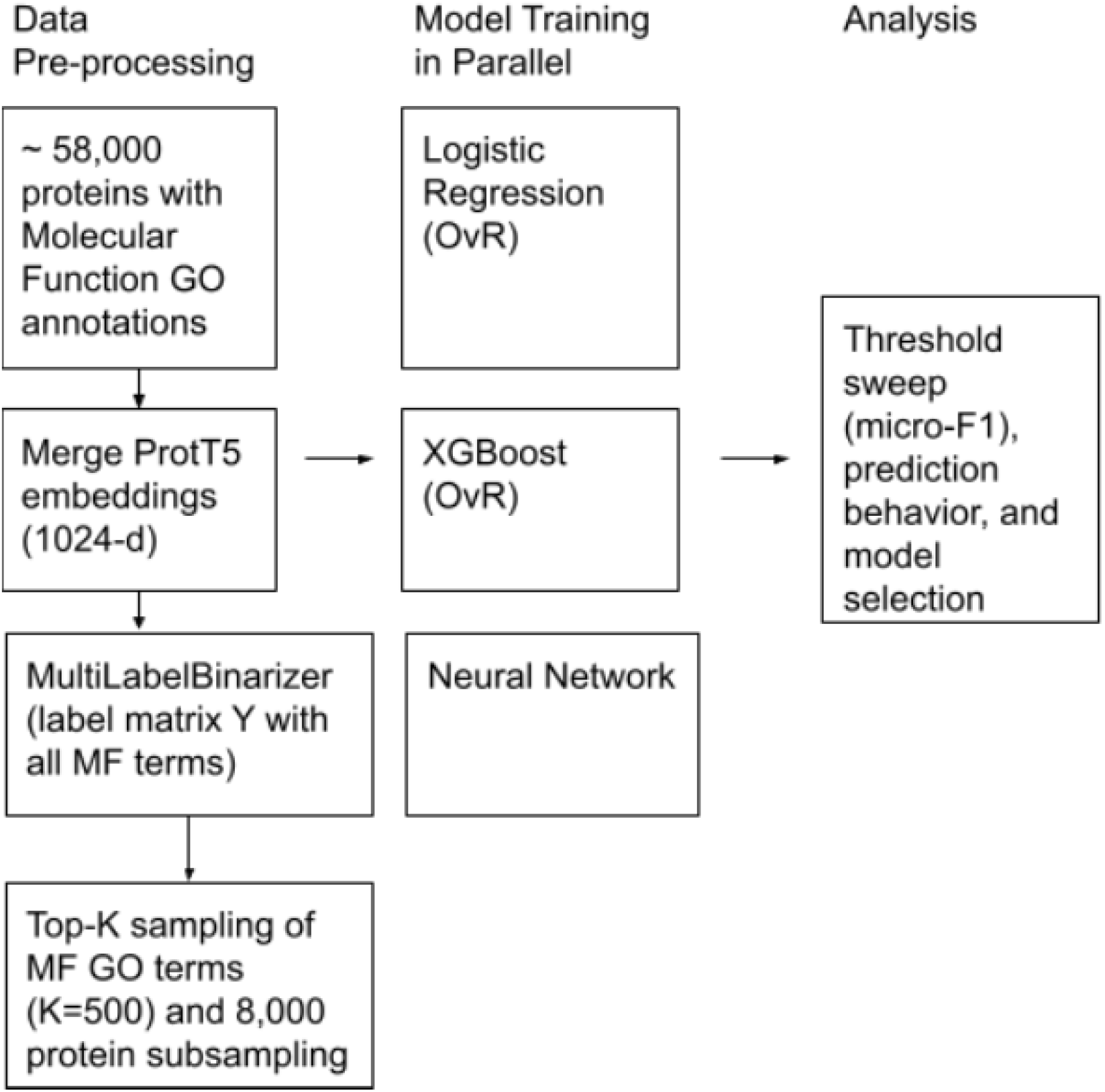
ML workflow for multi-classification detailing data preprocessing using ProtT5 embeddings, MultiLabelBinarizer, and Top-K sampling techniques. The data was trained using logistic regression (OvR), XGBoost (OvR), or a neural network, and analyzed using micro-F1 and threshold sweeps for model selection.

**Figure 2:**
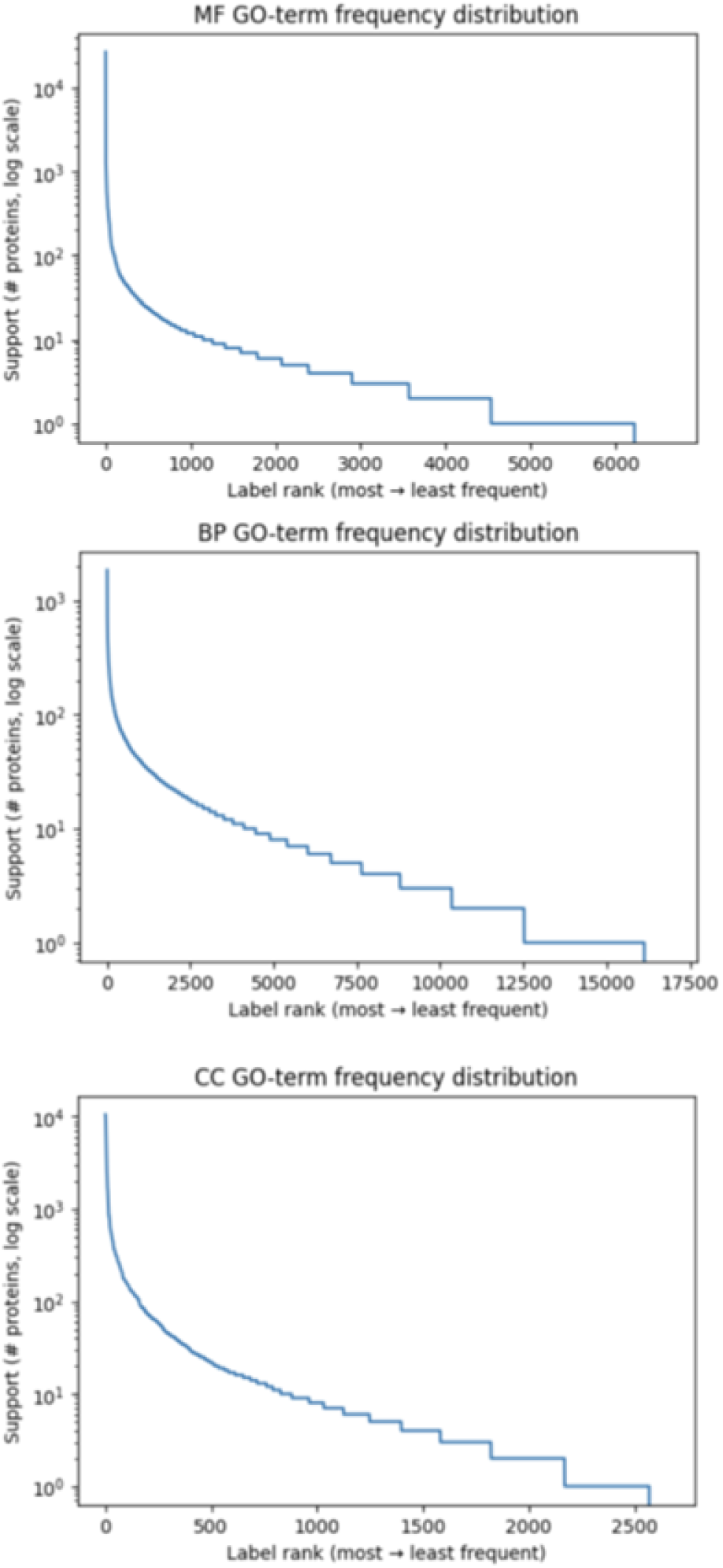
Ontology-specific GO-term frequency distributions for MF (top), BP (middle), and CC (bottom). All exhibit strong long-tail behavior, supporting the use of top-K sampling for stable model training.

**Figure 3:**
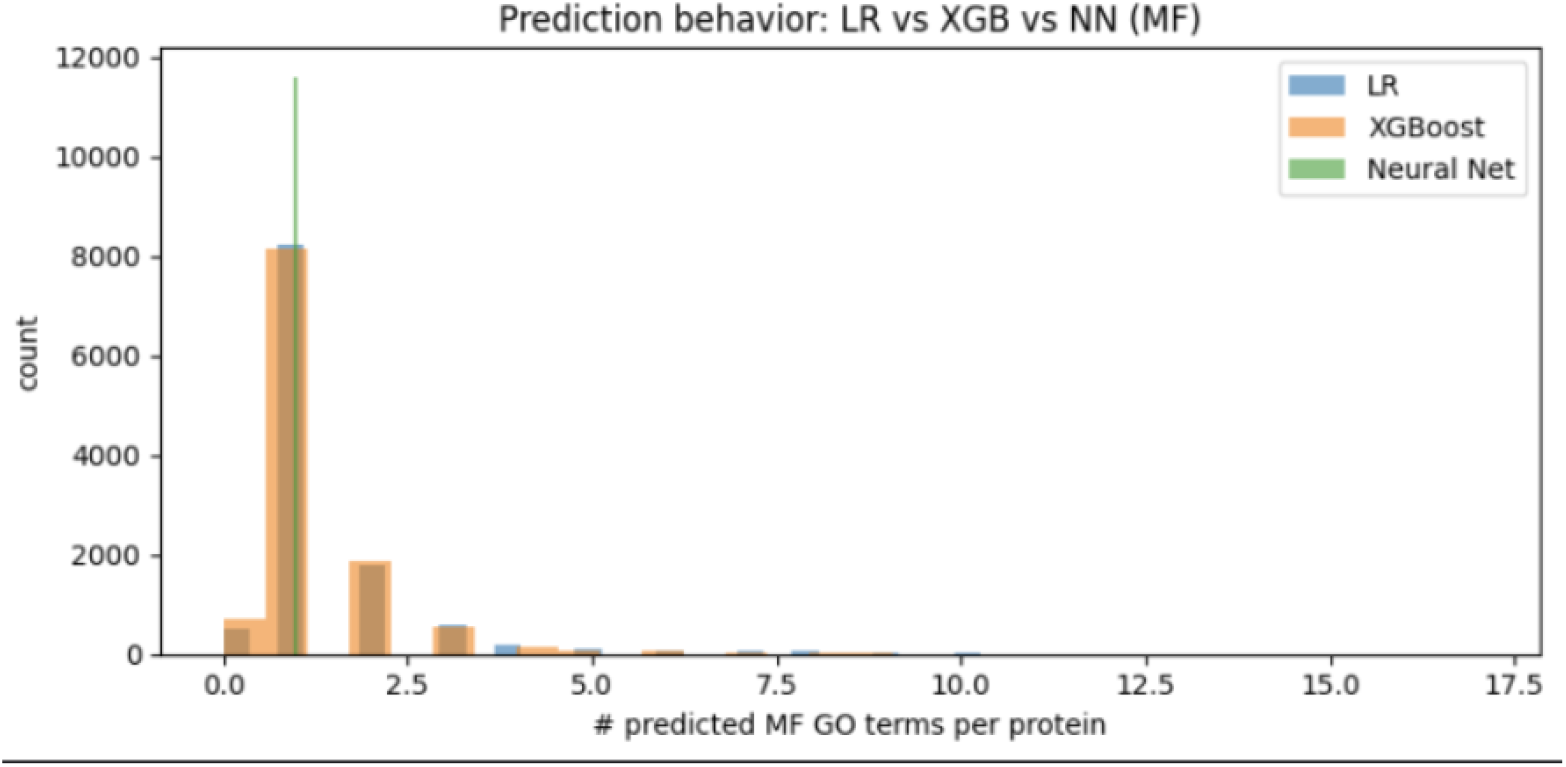
Distribution of predicted MF GO terms per protein for logistic regression, XGBoost, and neural network models at their respective operating thresholds. The neural network exhibits underprediction, while LR and XGBoost produce similar annotation volumes.

Three modeling approaches were compared. First, a One-vs-Rest logistic regression classifier [9] was trained using scikit-learn using a linear solver, maximum 200 iterations, and binary cross-entropy loss to assess whether non-linear methods offer measurable improvement.

Next, XGBoost (One-vs-Rest), an ensemble gradient boosting approach [10], was implemented. The following parameters were defined: binary logistic objective, histogram tree method, max depth = 5, learning rate = 0.1, and subsampling and column subsampling = 0.8. Predicted probabilities from individual classifiers were concatenated to produce a full multi-label probability matrix.

Additionally, a neural network was implemented using PyTorch, with the following parameters: input layer (1024 → 512), ReLU activation, dropout = 0.3, output layer (512 → 500), and sigmoid output activation. The model was trained for 10 epochs using Adam optimization and binary cross-entropy loss.

For each model, prediction thresholds in the range of 0.05–0.30 were evaluated. The threshold that maximized the micro-averaged F1 score [13] on the validation set was selected. Additionally, the number of predicted MF terms per protein, fraction of proteins that received no predictions, and performance stratified by GO term frequency (rare or common terms) was examined.

### 2.3 Ontology-specific models

Following preliminary model selection on the MF ontology, XGBoost (One-vs-Rest) was selected as the primary classifier for training ontology-specific predictors for BP and CC proteins. Since MF, BP, and CC have different semantics, label counts and sparsity profiles, training jointly would bias the model towards one specific ontology, and so three separate models were trained.

Since CAFA evaluates ontologies separately, this enables ontology-specific hyperparameters and thresholds, simplifying interpretation and deployment.

GO annotations were loaded from the CAFA-6 training file and grouped by ontology. ProtT5 embeddings were loaded from the precomputed embedding dataframe and inner-joined with BP-labeled proteins using EntryID. The data was binarized into a multi-hot label matrix using Multi-LabelBinarizer, producing feature matrix *X* ∈ℝ^*n*×1024^ and label matrix *Y* ∈ {0, 1}^*n×L*^, where *L* is the number of distinct BP terms prior to filtering.

Proteins were split into train and validation sets using an 80/20 split with a fixed random seed to ensure reproducibility.

Since all ontology labels exhibit an extreme long-tail distribution with very sparse terms, training was restricted to the Top-K most frequent terms. Since the GO labels follow a power law and long-tail distribution, support eventually dropped to tens of proteins per term, indicating too much sparsity to learn stable decision boundaries. Thus, *K* was chosen at the inflection region of the log-scale frequency curve where support transitions from steep decay to a more gradual tail (MF: *K* = 500; BP and CC analogous ranges), balancing label coverage and statistical support.

### 2.4 Test set inference

For test set inference, the trained ontology-specific XGBoost One-vs-Rest models for MF, BP, and CC proteins were loaded along with their corresponding metadata, including selected label indices and GO term mappings.

Protein embeddings for the CAFA-6 test set were generated using the same pretrained ProtT5 embedding pipeline applied to the training data.

Test protein identifiers were parsed from the CAFA-6 test FASTA file, and strict alignment between protein order and embedding vectors was verified prior to inference.

For each ontology, predicted probabilities were obtained using the trained OvR XGBoost classifiers, producing independent probability matrices for MF, BP, and CC.

Predictions were filtered using a global probability cutoff of *p* ≥10^*−*4^ to retain low-confidence but potentially informative GO term assignments while also ensuring computational speed.

All retained predictions were written to a single submission file containing protein identifiers, GO term identifiers, and predicted probabilities.

## 3 Results

### 3.1 Preliminary Model Comparison on Molecular Function

Logistic regression (LR), XGBoost (OvR), and a feedforward neural network were compared on a held-out MF validation subset. LR and XGBoost achieved similar peak micro-averaged F1 scores at comparable operating thresholds (≈ 0.15), whereas the neural network required a substantially higher threshold (≈ 0.55) and achieved lower peak performance, indicating poorer threshold behavior and score distribution.

Analysis of prediction behavior showed that LR and XGBoost produced highly similar numbers of predicted MF terms per protein, with substantial overlap in their prediction-count distributions. In contrast, the neural network exhibited near-degenerate behavior, predicting approximately one MF term per protein for most sequences, consistent with systematic underprediction.

Based on its slightly higher peak performance and more stable prediction behavior, XGBoost was selected for downstream ontology-specific modeling.

### 3.2 Molecular Function Model

Using the trained MF XGBoost One-vs-Rest model, predicted probabilities were generated on the validation set for the Top-500 most frequent MF GO terms. To understand how prediction behavior varies with the probability cutoff, thresholds in the range 0.05–0.30 were evaluated.

As the prediction threshold increased, the mean and median number of predicted MF terms per protein decreased monotonically (Figure 4). At low thresholds, the model produced broader annotations, while higher thresholds yielded more conservative predictions. Notably, the median number of predicted terms dropped from approximately two MF terms per protein at low thresholds to one MF term per protein beyond a threshold of ∼0.15.

**Figure 4:**
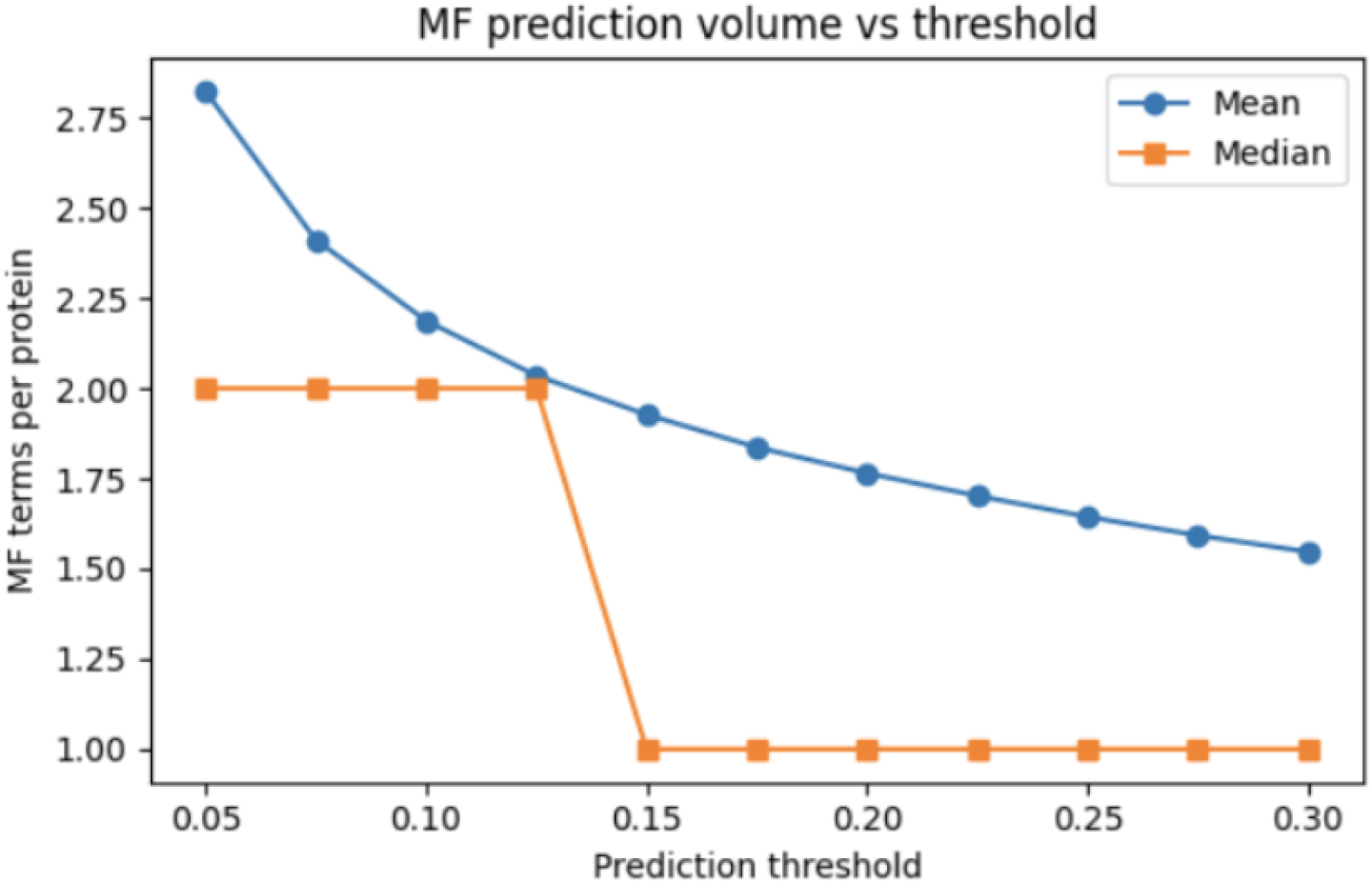
Mean and median number of predicted MF GO terms per protein as a function of prediction threshold.

In parallel, the fraction of proteins receiving no MF predictions increased with threshold (Figure 5). At a threshold of 0.05, fewer than 2% of proteins had zero predictions, whereas this fraction increased to approximately 14% at a threshold of 0.30, reflecting the trade-off between coverage and precision.

**Figure 5:**
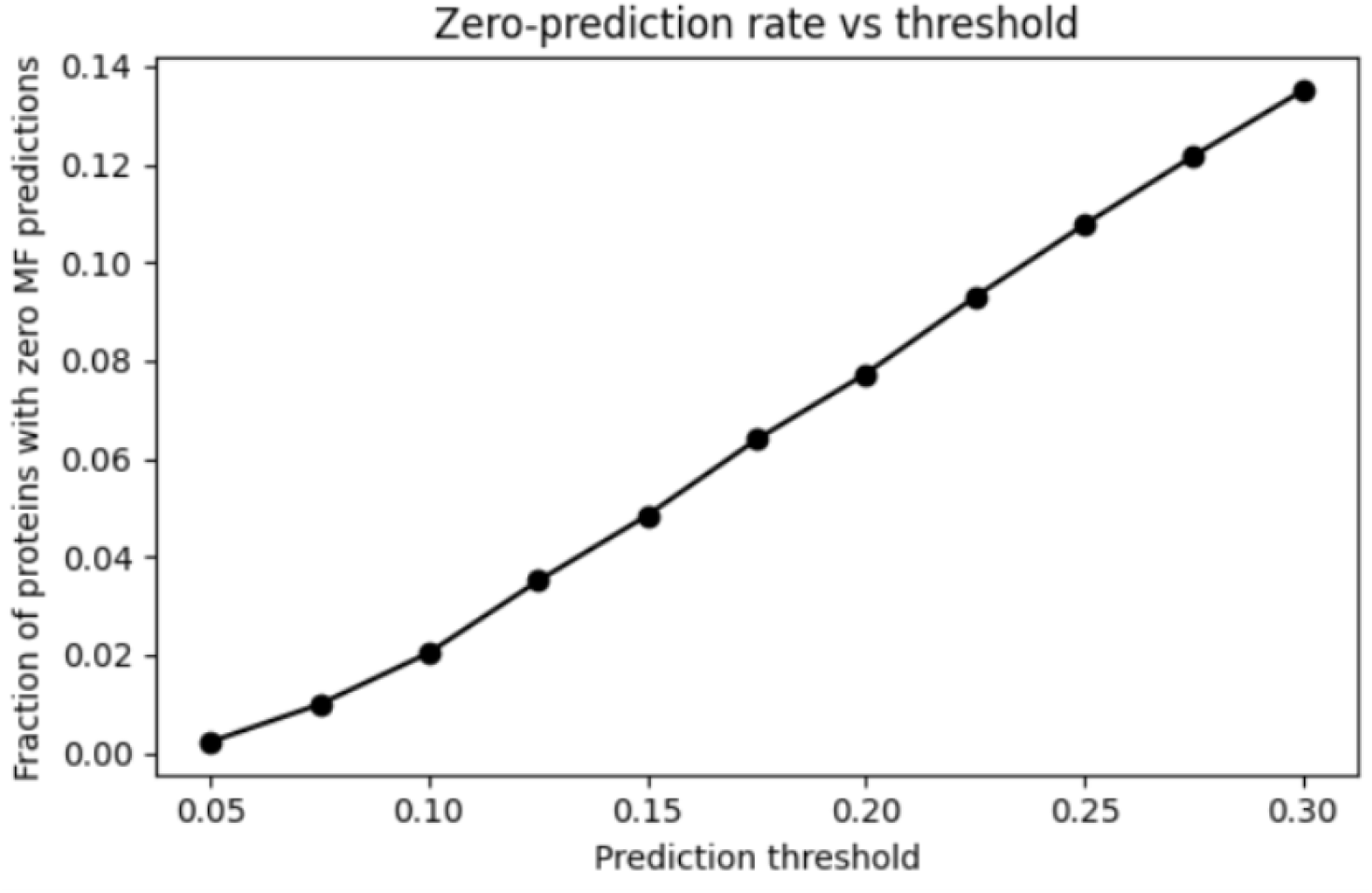
Fraction of proteins receiving zero MF predictions as a function of threshold.

### 3.3 Micro-F1 Optimization and Operating Point Selection

Micro-averaged F1 score was computed across thresholds to identify an operating point balancing precision and recall. Micro-F1 increased steadily with threshold up to approximately 0.20–0.25, after which performance plateaued and slightly declined (Figure 6). The optimal micro-F1 was achieved near this range, motivating the selection of a threshold in this region for further analysis.

**Figure 6:**
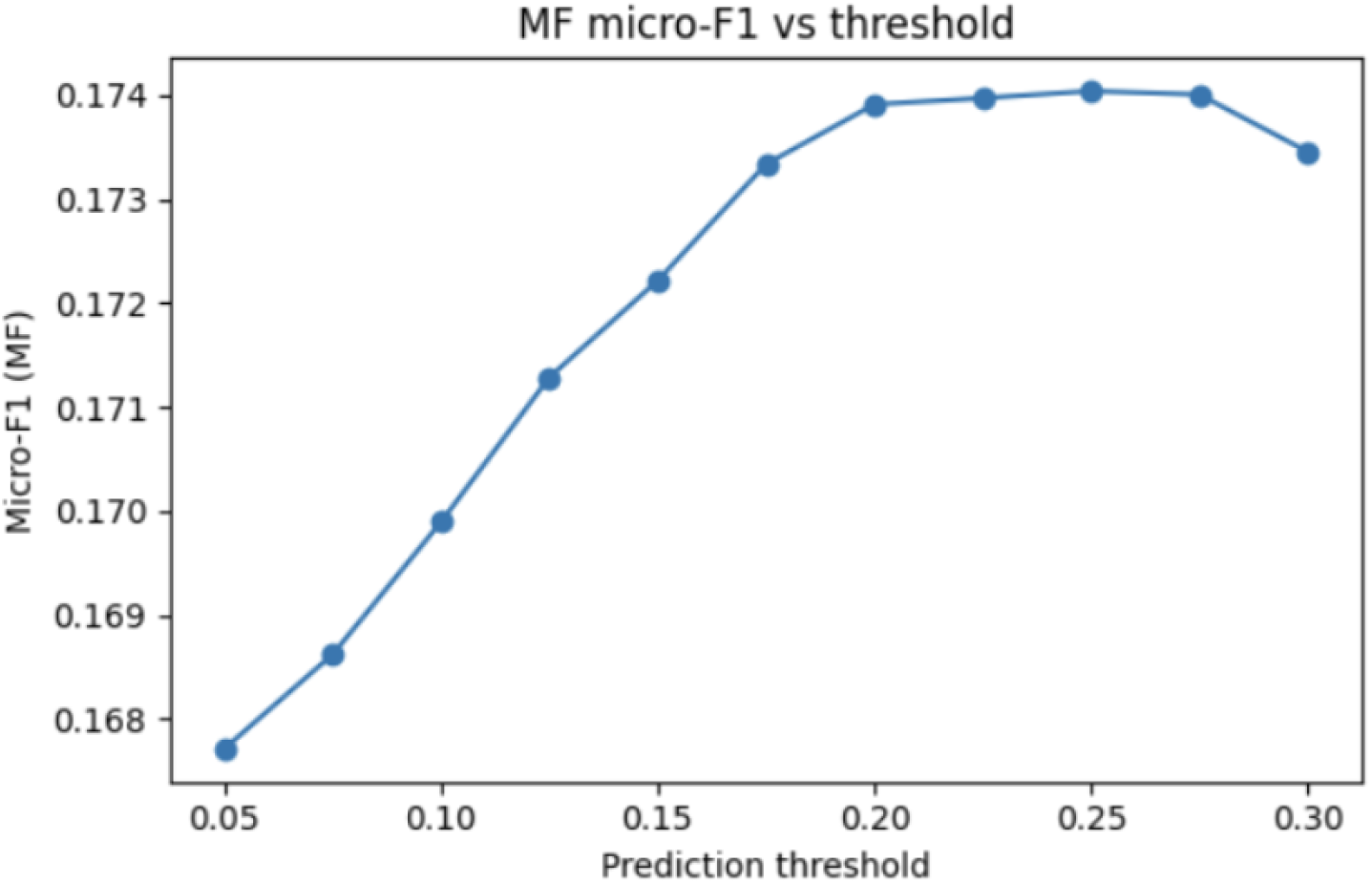
Micro-F1 score as a function of prediction threshold for MF ontology.

### 3.4 Agreement Between Predicted and True MF Annotation Counts

At a representative operating threshold (*t* = 0.20), the relationship between the number of true MF annotations per protein and the number of predicted MF annotations was examined (Figure 7). While the model tended to underpredict MF terms for proteins with large numbers of annotations, a positive correlation was observed overall, indicating that proteins with richer MF annotations generally received more predicted terms.

**Figure 7:**
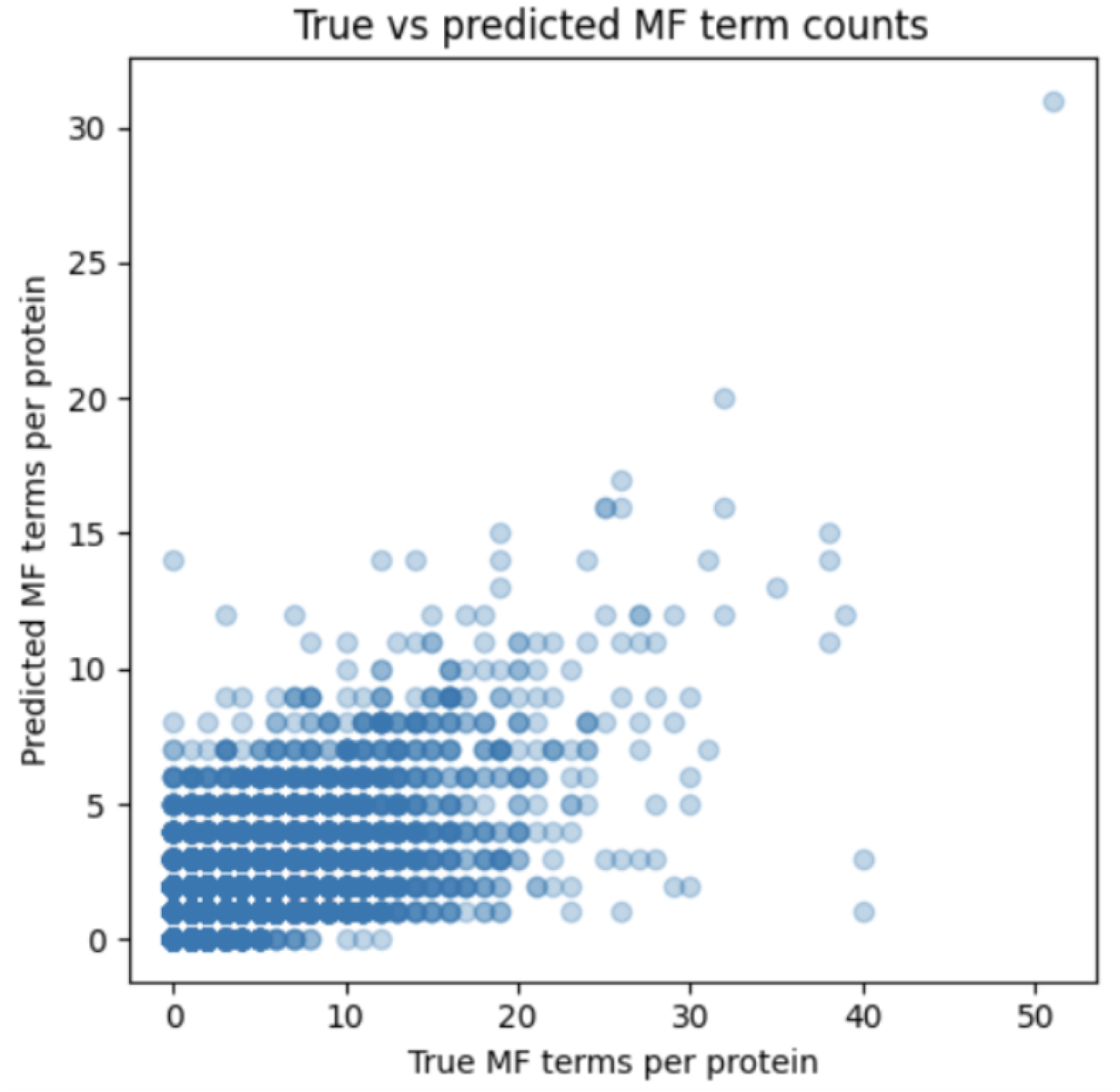
Relationship between true and predicted MF term counts per protein at the selected threshold.

### 3.5 Error Profile and Label Difficulty

An aggregate confusion analysis at the selected threshold showed that the majority of errors were false negatives, reflecting the inherent difficulty of predicting sparse MF annotations. True positives accounted for a minority of all positive events, while false positives remained relatively constrained compared to false negatives.

Per-label recall analysis revealed that low-support MF terms were substantially more difficult to predict, with many rare terms exhibiting near-zero recall. This behavior is consistent with the long-tail label distribution and further supports the Top-K restriction strategy used during training.

### 3.6 Biological Process (BP) Results

The BP XGBoost One-vs-Rest model was evaluated on the validation set using the Top-1000 most frequent BP GO terms. As expected given the substantially larger label space and higher annotation density of BP compared to MF, the model produced a much larger number of predicted terms per protein at low probability thresholds (Figure 8).

**Figure 8:**
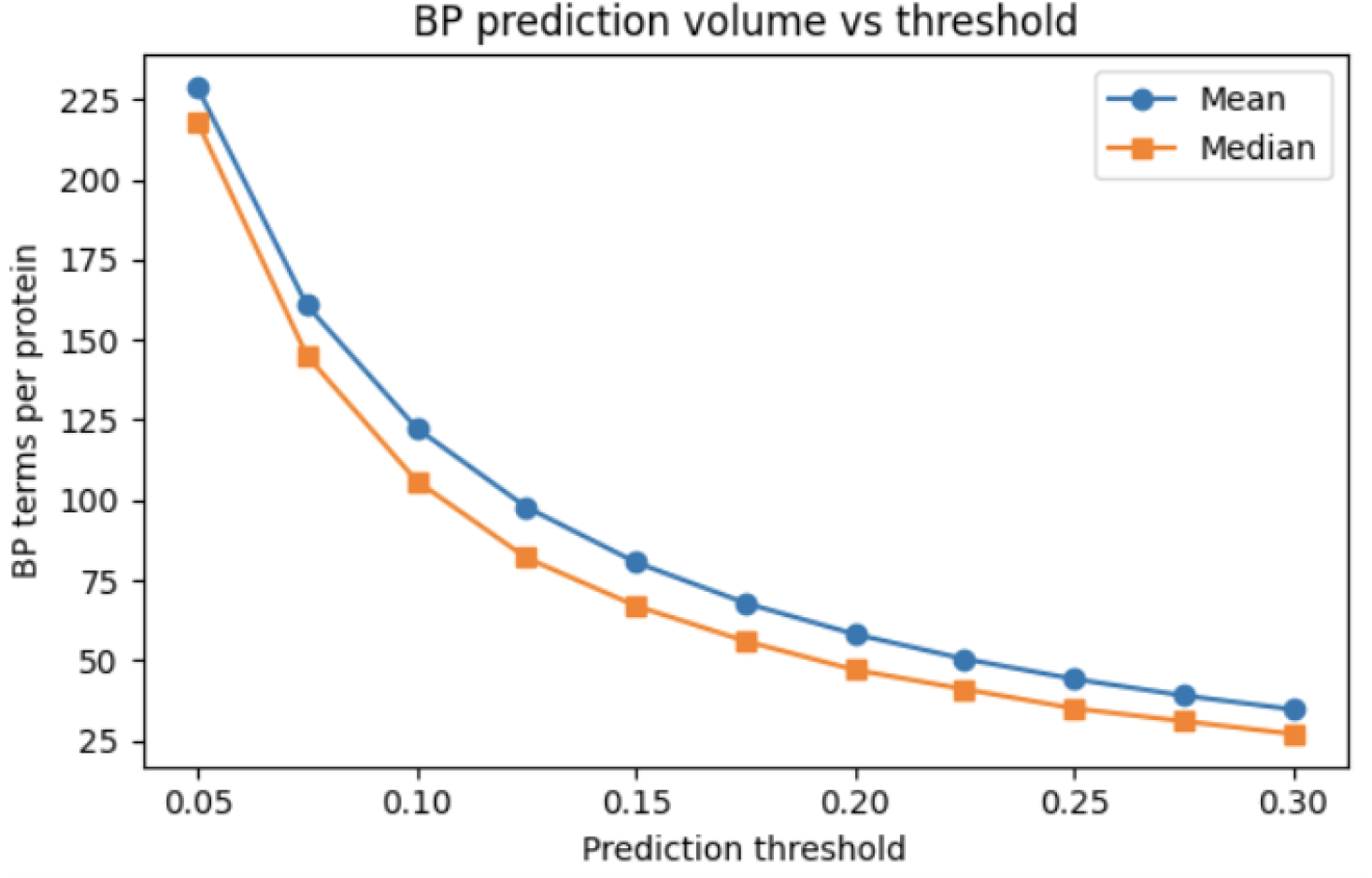
Mean and median number of predicted BP terms per protein across thresholds.

As the prediction threshold increased from 0.05 to 0.30, the mean and median number of predicted BP terms per protein decreased smoothly, without abrupt collapse. At lower thresholds, the model assigned broad BP annotations, while higher thresholds yielded more conservative predictions. Importantly, the fraction of proteins receiving zero BP predictions remained near zero across most thresholds, increasing only at the most stringent cutoffs (Figure 9), indicating that the model maintains broad coverage even under conservative operating points.

**Figure 9:**
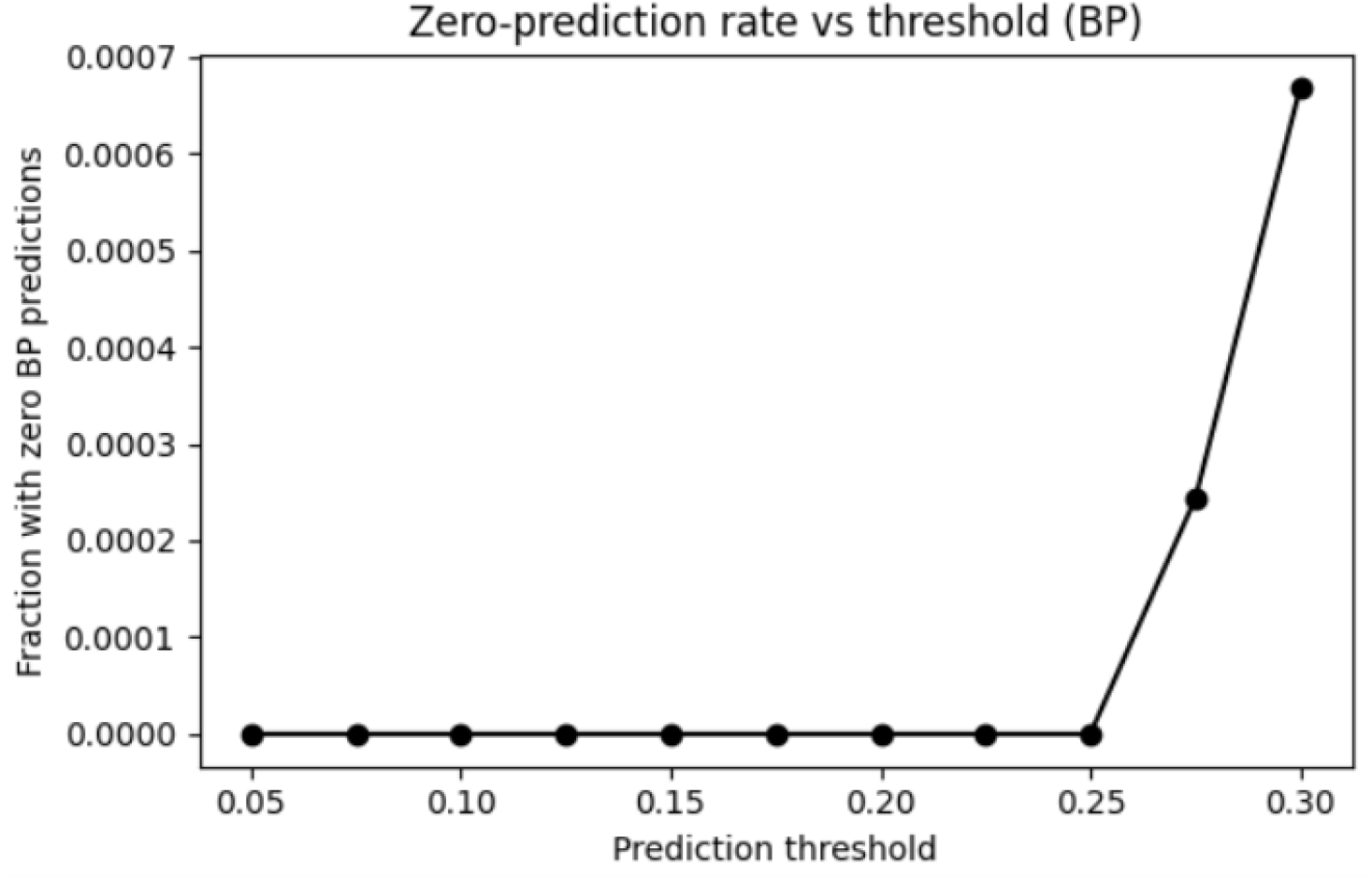
Fraction of proteins receiving zero BP predictions across thresholds.

#### Micro-F1 behavior and operating point considerations (BP)

Micro-averaged F1 scores for BP were uniformly low across thresholds (Figure 10), reflecting the extreme label sparsity and semantic breadth of the BP ontology. However, the micro-F1 curve remained smooth and stable, without sharp discontinuities, suggesting consistent model behavior rather than threshold-sensitive instability.

**Figure 10:**
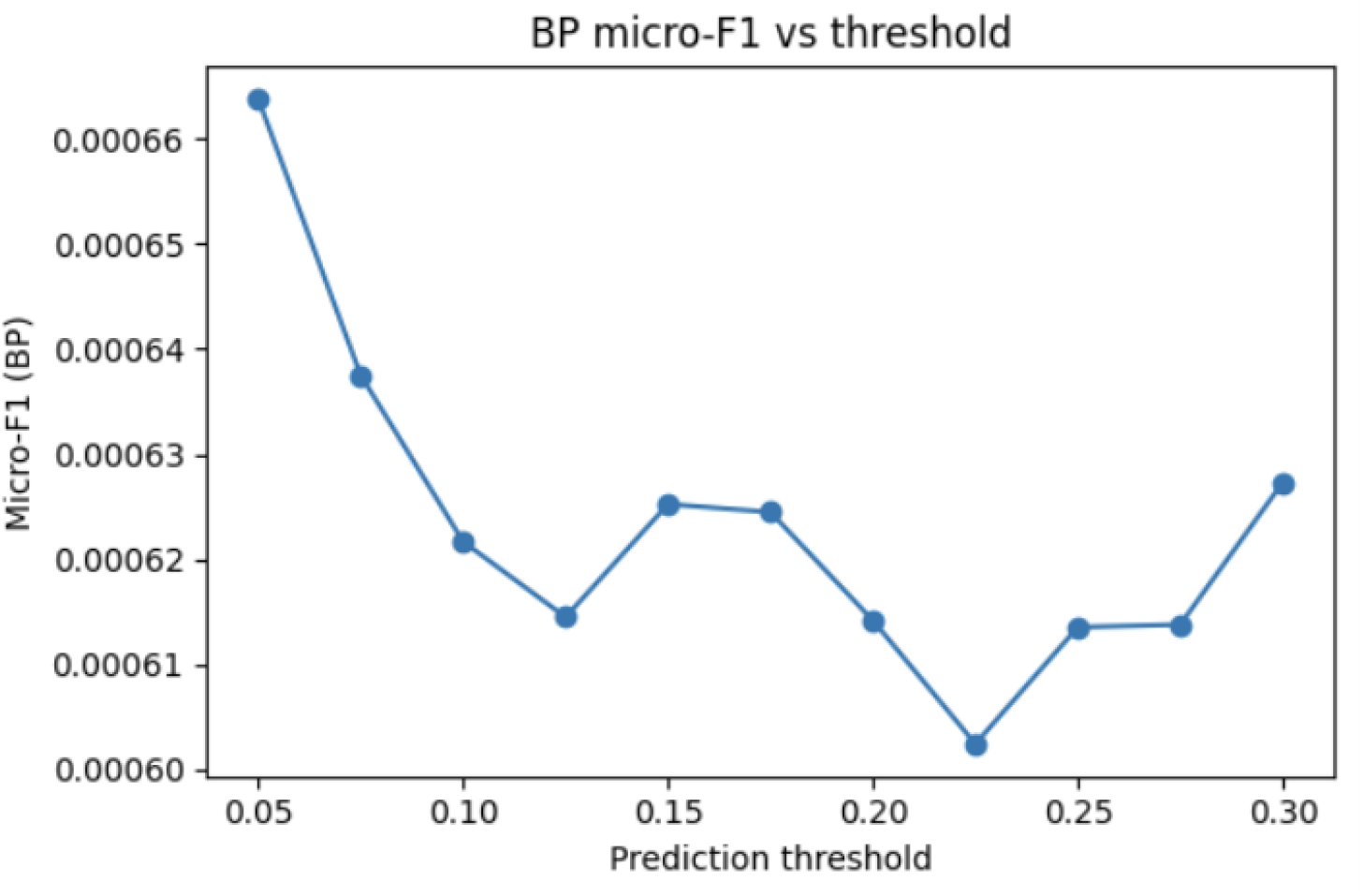
Micro-F1 score across thresholds for BP ontology.

Rather than optimizing aggressively for a single threshold, an operating range was selected based on prediction stability and biological plausibility, prioritizing reasonable annotation volume and minimal zero-prediction behavior.

#### Agreement between predicted and true BP annotation counts

At a representative operating threshold, the relationship between true and predicted BP term counts per protein was examined (Figure 11). The model tended to overpredict BP terms for sparsely annotated proteins, while partially tracking proteins with richer BP annotations.

**Figure 11:**
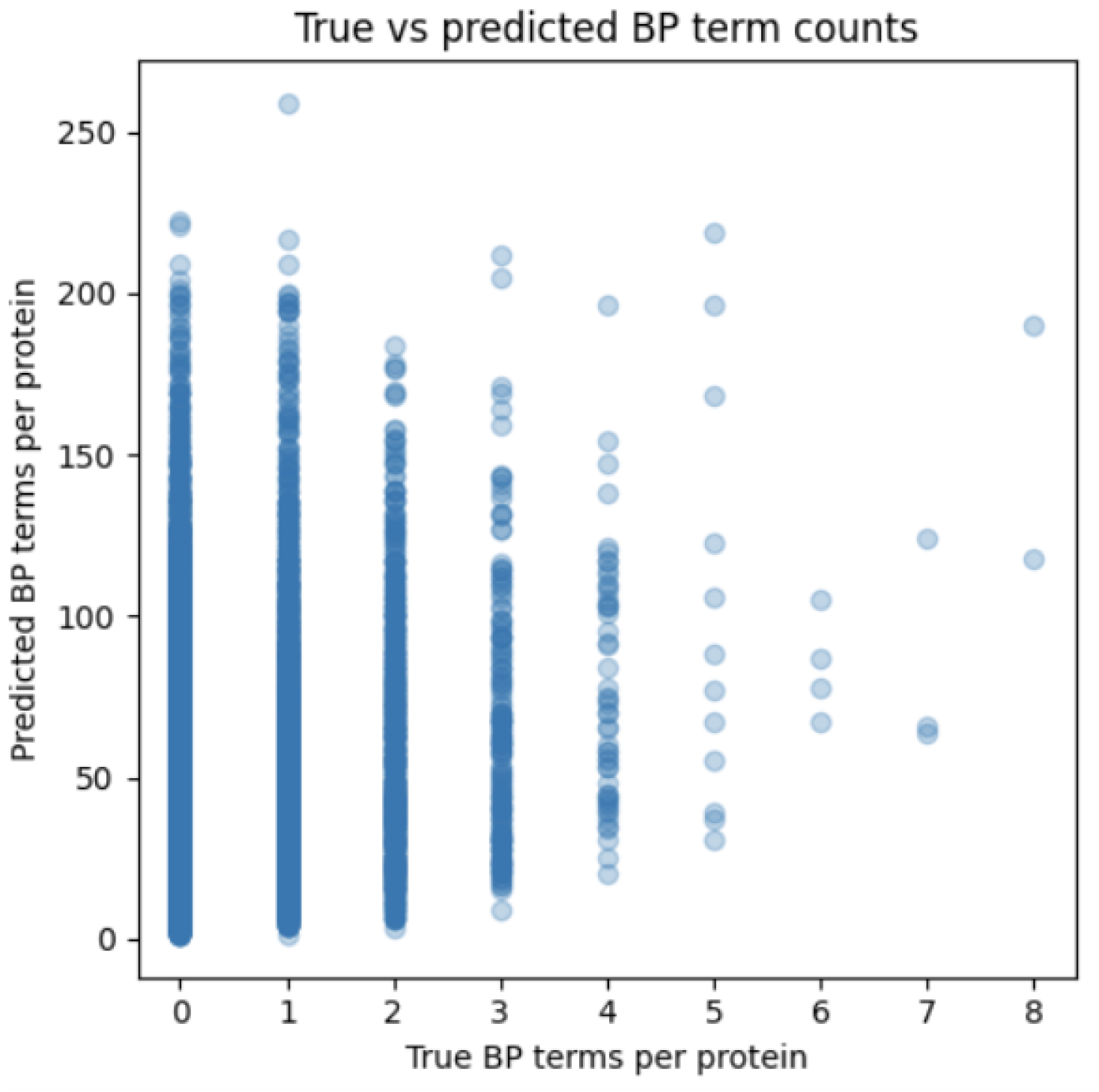
True vs predicted BP term counts per protein.

Perfect correspondence is not expected in BP due to the hierarchical and overlapping nature of biological process terms, but the observed trend indicates that the model captures coarse-grained annotation density.

### 3.7 Label Difficulty and Long-Tail Effects

Per-label recall analysis revealed that rare BP terms were particularly difficult to predict, with many low-support terms exhibiting zero recall. This behavior is consistent with the long-tail distribution of BP labels and further motivates the Top-K restriction used during training.

### 3.8 Cellular Component (CC) Results

#### Prediction volume and threshold sensitivity

The CC XGBoost model was trained on the Top-300 most frequent CC terms and evaluated on the validation set. Compared to MF and BP, CC predictions exhibited substantially higher annotation density per protein, particularly at lower thresholds (Figure 12). At a threshold of 0.05, the model predicted dozens of CC terms per protein on average, reflecting the finer-grained localization structure of CC annotations.

**Figure 12:**
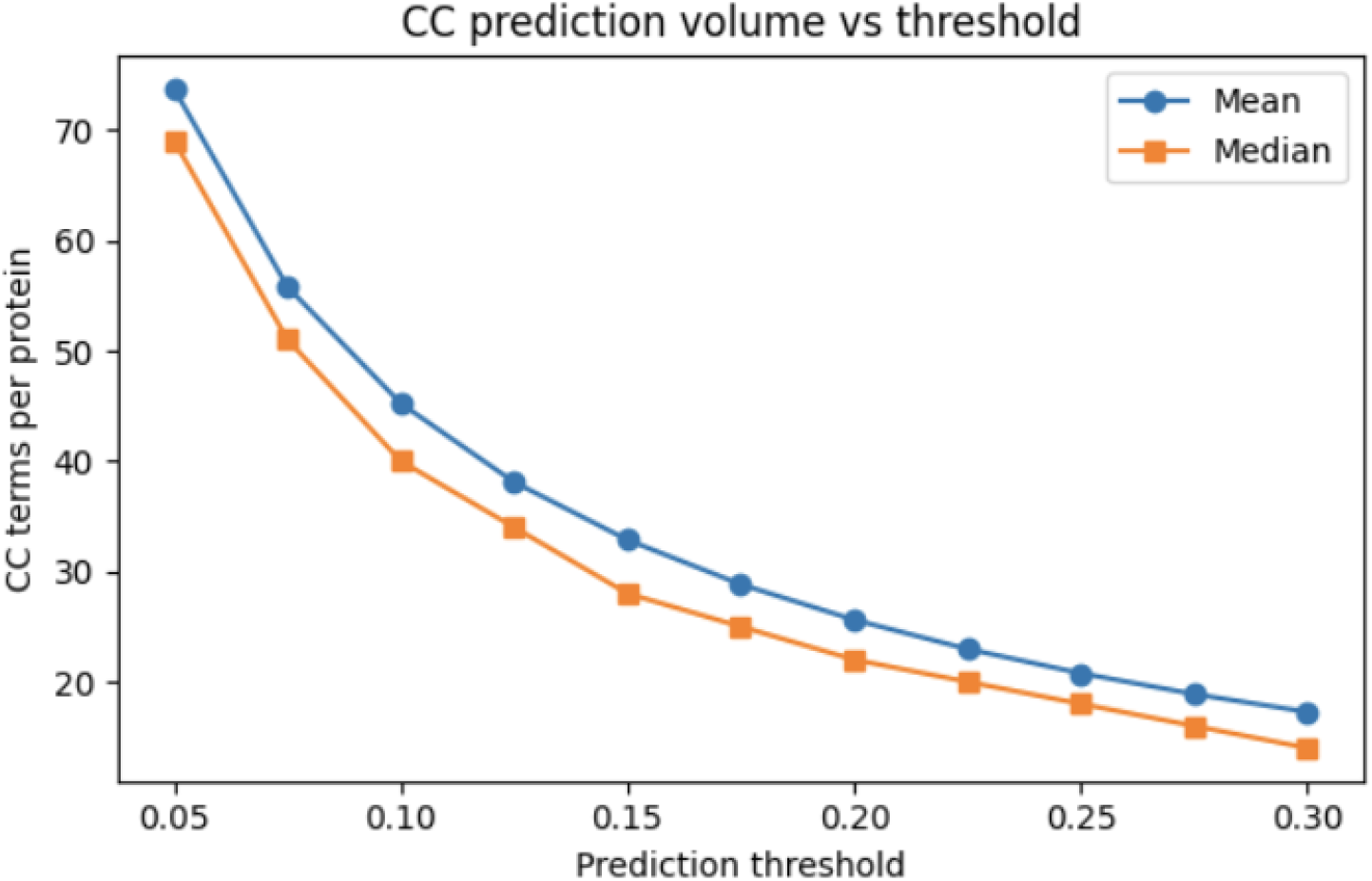
Mean and median number of predicted CC terms per protein across thresholds.

As the threshold increased, the number of predicted CC terms per protein decreased smoothly, indicating stable probability calibration rather than abrupt pruning.

#### Micro-F1 trends and interpretation

Absolute micro-F1 values for CC were low across all thresholds (Figure 13), which is expected given the highly overlapping and spatially granular nature of CC labels. Importantly, the micro-F1 curve increased monotonically with threshold, demonstrating consistent trade-offs between precision and recall rather than noisy or unstable behavior.

**Figure 13:**
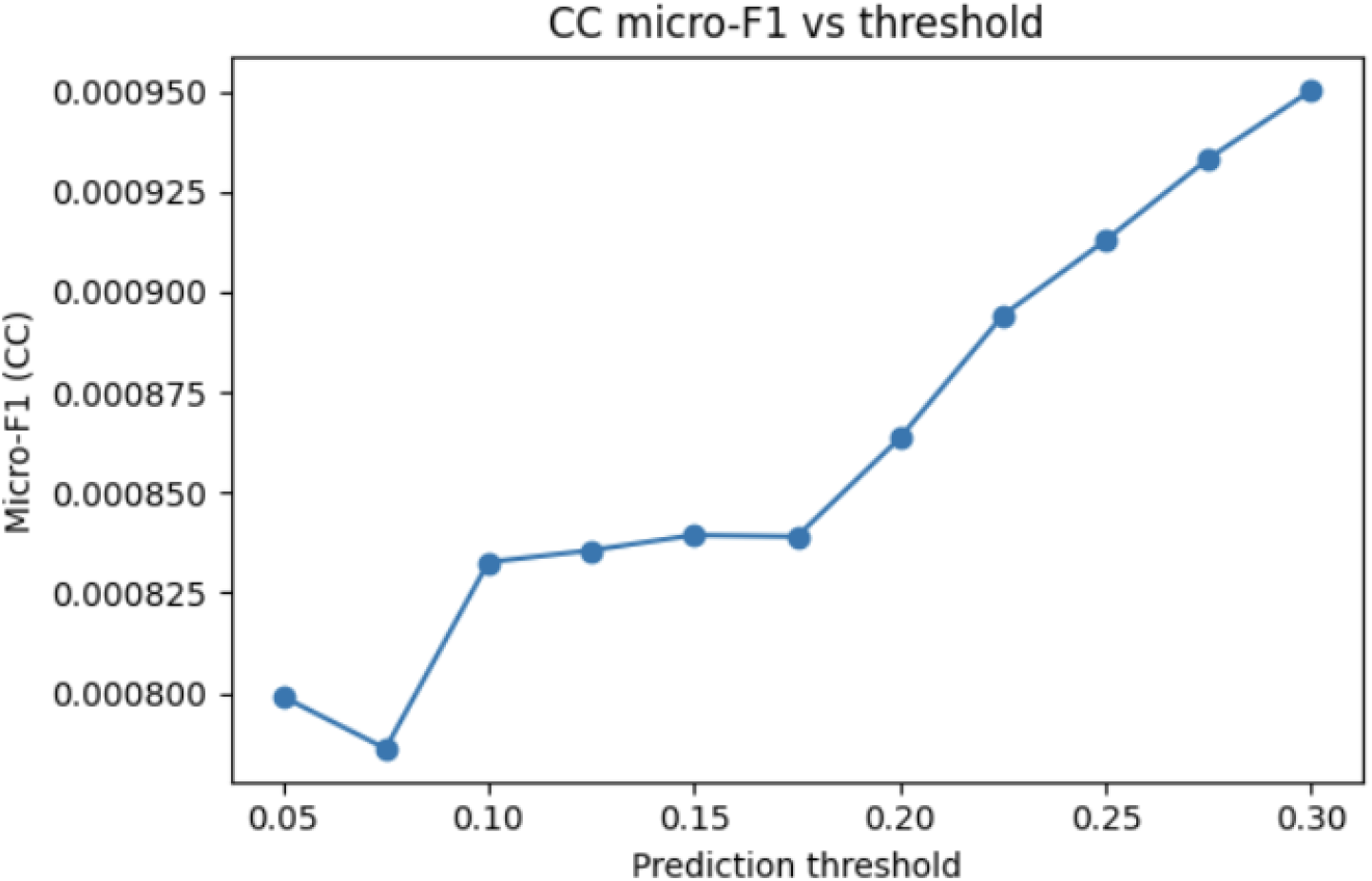
Micro-F1 score across thresholds for CC ontology.

Micro-F1 values for CC should therefore be interpreted primarily as a relative diagnostic, rather than as an absolute performance measure.

#### True vs predicted CC annotation counts

Comparison of true and predicted CC term counts per protein revealed systematic overprediction, particularly for proteins with few annotated CC terms (Figure 14). This reflects both annotation incompleteness and the tendency of sequence-based models to predict multiple plausible cellular localizations.

**Figure 14:**
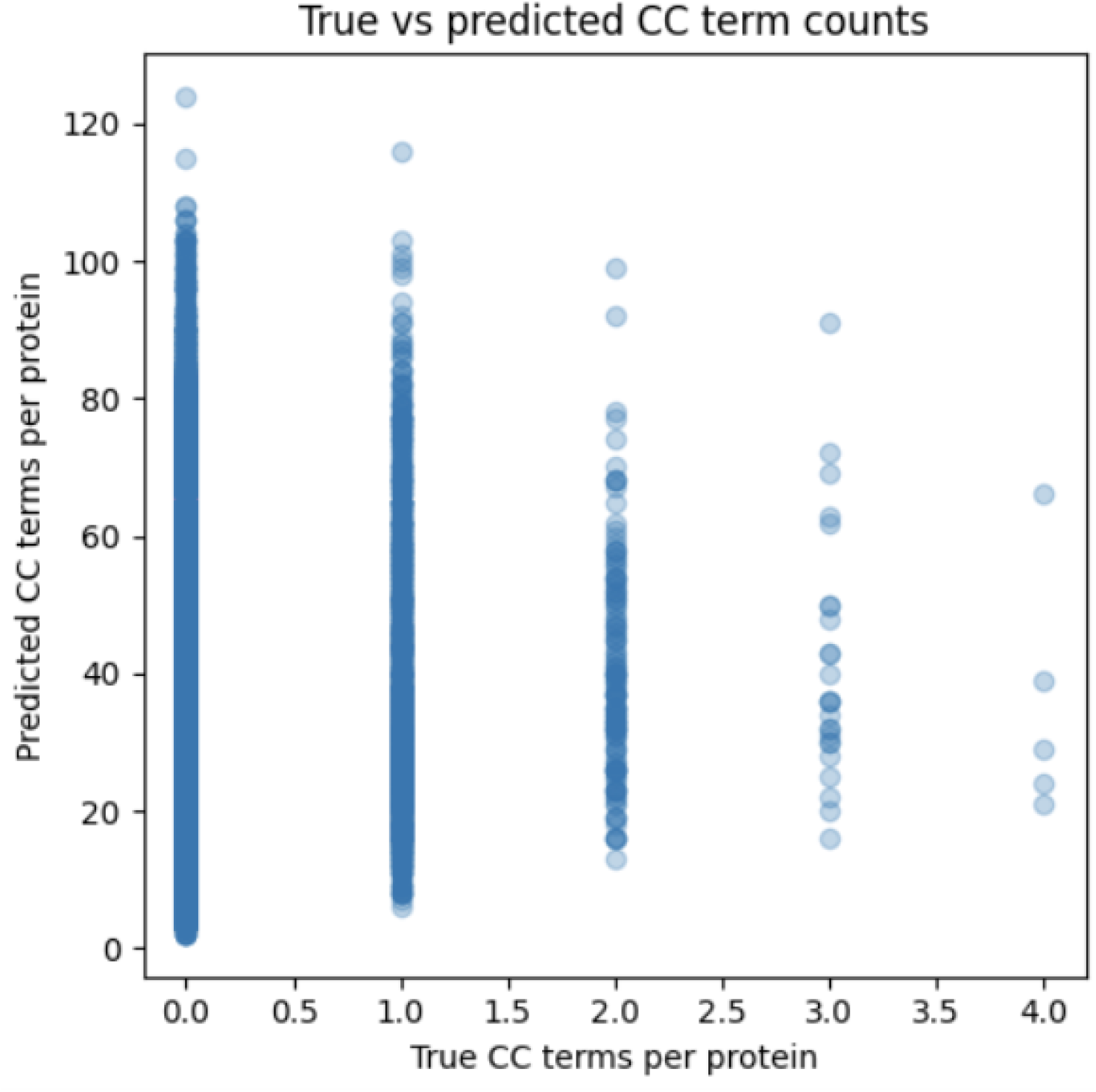
Relationship between true and predicted CC term counts per protein at a representative threshold.

Despite this overprediction, proteins with more annotated CC terms generally received higher numbers of predicted terms, indicating partial alignment between predicted and true annotation complexity.

#### Rare label performance

Per-label recall analysis showed that CC terms with very low support (often single-digit counts) were rarely recovered. These terms typically correspond to highly specific subcellular structures or rare compartments, which are difficult to infer reliably from sequence alone.

### 3.9 Summary Across Ontologies

Across MF, BP, and CC, the XGBoost-based models exhibited stable, interpretable threshold-dependent behavior, with smooth trade-offs between prediction coverage and confidence. While absolute performance varied substantially across ontologies due to differences in label density and semantic scope, the results demonstrate that pretrained transformer embeddings combined with gradient-boosted classifiers provide a consistent and scalable framework for ontology-specific protein function prediction.

## 4 Conclusion

In this study, we evaluated a practical pipeline for large-scale protein function prediction using pretrained transformer embeddings and supervised multi-label classifiers within the CAFA-6 setting. By leveraging ProtT5-derived sequence embeddings, we avoided handcrafted features and alignment-based methods while retaining rich contextual information about protein sequences.

Our analysis highlighted the severe long-tail structure of Gene Ontology annotations and motivated the use of Top-K label filtering to ensure stable model training.

Across preliminary comparisons, gradient-boosted decision trees consistently provided the best balance between predictive performance and stable prediction behavior, outperforming both linear models and a shallow neural network on the Molecular Function validation task. Training separate ontology-specific models further simplified optimization and reflected fundamental differences in label distributions across Molecular Function, Biological Process, and Cellular Component annotations.

While overall performance remains constrained by label sparsity and incomplete annotations, this work demonstrates that pretrained protein language model embeddings combined with carefully chosen classical classifiers form a strong and interpretable baseline for protein function prediction. Future work may include incorporating Gene Ontology hierarchy information during training, improved modeling of rare labels, ensembling across complementary classifiers, and integrating structural information such as AlphaFold-derived representations to better capture functions that are difficult to infer from sequence alone.

## Author Contributions

Jett Appel designed the study, implemented the modeling pipeline, conducted all experiments, performed analysis, and wrote the manuscript. Nathan Butcher contributed to embedding generation and provided feedback on the project direction.

